# *Mycobacterium smegmatis* NucS-promoted DNA mismatch repair involves limited resection by a 5’-3’ exonuclease and is independent of homologous recombination and NHEJ

**DOI:** 10.1101/2023.11.26.568737

**Authors:** Iris V. Rivera-Flores, Emily X. Wang, Kenan C. Murphy

**Affiliations:** Department of Microbiology and Physiological Systems University of Massachusetts Chan Medical School Worcester, MA 01605

## Abstract

The MutSL mismatch repair (MMR) system in bacteria and eukaryotes correct mismatches made at the replication fork to maintain genome stability. A novel MMR system is represented by the EndoMS/NucS endonuclease from actinobacterium *Corynebacterium glutamicu*m, which recognizes mismatched substrates in vitro and creates dsDNA breaks at the mismatch. In this report, a genetic analysis shows that an *M. smegmatis* Δ*nucS* strain could be complemented by expression of wild type NucS protein, but not by one deleted of its last five amino acids, a region predicted to be critical for binding to the β-clamp at the replication fork. Oligo-recombineering was then leveraged to deliver defined mismatches to a defective hygromycin resistance gene on the *M. smegmatis* chromosome. We find that NucS recognizes and repairs G-G, G-T, and T-T mismatches in vivo, consistent with the recognition of these same mismatches in C. *glutamicum* in vitro, as well as mutation accumulation studies done in *M. smegmatis*. Finally, an assay that employs an oligo that promotes the generation of two mismatches in close proximity on the chromosome shows that a NucS-promoted cut is processed by a 5’–3’ exonuclease and that NucS-promoted MMR is independent of both homologous recombination and non-homologous end-joining.

## Introduction

High fidelity of DNA replication in all organisms is of paramount importance to minimize mutation accumulation and maintain genome stability. Most cells correct the misincorporation of the wrong nucleotide during DNA replication by two distinct mechanisms. The first line of defense is provided by the proofreading functions of DNA polymerases that immediately detect and exonucleolytically remove an incorrect nucleotide in the nascent DNA strand (1–3). Should this system fail, however, DNA mismatch repair (MMR) systems recognize mismatched base pairs in the wake of the replication fork, identify which base of the mismatch is the incorrect one, and processes its removal (4,5). Resynthesis of the exposed template and religation to the undamaged preexisting DNA strand completes the repair process. The canonical MMR pathway is represented by the MutSL system, which was first identified in *E. coli* (6,7). The *E. coli* MutS protein recognizes the mismatched bases and together with MutL promotes a platform for nicking of the newly synthesized strand by MutH endonuclease, governed by the hemi-methylation status of that strand. In most bacteria, however, *mutH* is not present and a latent MutL endonuclease is responsible for cutting the replicative strand (for reviews, see (8–11)). Homologous MutSL systems are found ubiquitously in both prokaryotic and eukaryotic organisms, including humans (5). Mutations of these MMR genes lead to hyper-mutagenic genetic backgrounds, increased rates of homeologous recombination, and cancer.

Most archaeal and many actinobacterial genomes (including mycobacteria) are devoid of the canonical MutSL MMR system. It was believed for many years that perhaps these organisms did not possess an MMR system.

However, these species do not show higher rates of mutagenesis relative to bacteria that contain MutSL systems, suggesting that DNA replication fidelity in these bacteria relies solely on an efficient polymerase proofreading mechanism, or that an alternative MMR system exists in these species. The first hint of such a repair system came from the studies of Ishino *et al* (12) that described an endonuclease from the hyperthermophilic archaeon *Pyrococcus furiosus* that specifically recognizes and cuts mismatched dsDNA, which they called EndoMS (endonuclease specific for mismatch-specific DNA). EndoMS is homologous to a protein previously isolated from *Pyrococcus abyssi*, that was identified as a nuclease specific for ssDNA, called NucS (13), which is the name used for this protein hereafter.

Ichino *et al* (12) went on to study the NucS protein from *Thermococcus kodakarensis* (TKO), a more genetically tractable thermophilic archaeon. They found that the TKO NucS specifically binds to and cuts both strands of dsDNA species that contained G-T, G-G and T-T mismatches. Higher concentrations of the enzyme were required to cut dsDNA containing T-C and A-C mismatches, though such substrates failed to bind TKO NucS in an electrophoretic mobility shift assay. DNA substrates containing C-C, A-C and A-A showed no binding or cutting of mismatched substrates.

NucS proteins have no homology to any of the components of the mutSL systems, and thus was the first protein from an archaeon that suggested the existence of an alternative MMR system. Enzymatically, TKO NucS cuts on both sides of the mismatch leaving 5’-P and 3’-OH ends with 5’ protruding overhangs, reminiscent of the action of restriction enzymes. Indeed, the structure of the TKO NucS bound to 15-mer mismatched-containing dsDNA was solved by Nakae *et al* (14) showing that NucS protein to be remarkably similar to the structure of type II restriction enzymes, with the N-terminal domain promoting dimerization of NucS and the C-terminal domain binding to regions flanking the mismatched bases. The two bases of the mismatch were flipped out of the helix into binding sites within the protein, not unlike the recognition sequence of the Ecl18kl restriction enzyme (CCNGG), where the central base pair is flipped out in a manner like NucS acting on a mismatch. Unlike restriction enzymes, however, NucS does not have a recognition sequence in that it does not contact any bases at the cut site except for those contained within the mismatch. The recognition and cutting of mismatches by TKO NucS highlight the difference between this novel MMR system and that of MutSL, where MutS recognizes the mismatch, but a single-stranded DNA nick is made either by MutH directed to GATC sites on the unmethylated strand of the replication fork (*e.g.,* in *E. coli*), or in other bacteria and eukaryotes, by a β-clamp activation of a latent MutL endonuclease that nicks the newly synthesized strand. For recent updates on NucS endonuclease function and strand discrimination in MMR, see the following reviews (15,16).

Of particular interest is whether the novel NucS MMR system exhibited in the archaea described above is active as an anti-mutator in *Actinobacteria* where the gene is present, including the clinically relevant bacterium *Mycobacterium tuberculosis*. The question was addressed by two labs (17,18) that studied actinobacterium *Corynebacterium glutamicum* and found that deletion of the *nucS* homolog (NCgl-1168, hereafter called NucS_Cg_) resulted in a strain with a mutator phenotype. Purification of the NucS_Cg_ protein showed that it bound to DNA containing base pair mismatches, and that cutting of these substrates was activated by an interaction with the β-clamp from *C. glutamicum*, DnaN_Cg_. The mismatch recognition and cutting specificities of NucS_Cg_ were the same as those observed for TKO NucS and were consistent with the mutation spectrum of *C. glutamicum* in mutational accumulation (MA) studies done in the *nucS*-deficient background, *i.e.,* a higher levels of transitions observed relative to wild type (18).

This question of whether NucS from a mycobacteria species could bind mismatched DNA was addressed by Castañeda-Garcia *et al* (19) who examined thousands of transposon mutants in *Mycobacterium smegmatis* for high rates of spontaneous resistance to rifampicin. They identified MSMEG_4923 as a gene responsible for generating a mutator phenotype in *M. smegmatis*; this gene product has 27% sequence identity with NucS from *P. abyssi* and 73% sequence identity with NucS_Cg_. However, isolation and characterization of the *M. smegmatis* NucS protein showed that while it bound to ssDNA and not dsDNA (like its *P. abyssi* homolog), it did not demonstrate binding activity to mismatch containing dsDNA substrates in vitro. This result contrasts with those from the NucS proteins from *P. furiosus* and *T. kodakarensis* in vitro, raising the question of whether the mycobacterial NucS acts in a similar way to these proteins in vivo.

We have addressed this question here genetically by examining if *M. smegmatis* NucS requires an interaction with the β-clamp in vivo, as was seen previously with the NucS protein from *T. kodakarensis* in vitro. Complementation studies with plasmid-encoded wild type NucS and a derivative containing a C-terminal 5 amino acid truncation suggest an interaction with the *M. smegmatis* β-clamp is required for MMR activity in vivo. Secondly, we examined the mismatch specificity of *M. smegmatis* NucS in vivo and compared it to the mismatch specificity of *T. kodakarensis* and *C. glutamicum* reported in vitro. We used Che9 RecT-promoted oligo recombineering in *M. smegmatis* to directly deliver all possible types of base mismatches (except C-C) to a defective hygromycin-resistant (Hyg^R^) gene reporter construct integrated into the chromosome. We find that the specificity of *M. smegmatis* mismatch repair in vivo matches that found previously for *T. kodakarensis* in vitro, suggesting a commonality in the NucS-promoted MMR systems in these bacteria. Finally, we used oligo-mediated recombineering to deliver defined mismatches to the defective hygromycin-resistant target in the chromosome and find that a cut induced by NucS at a G-G mismatch is processed by a 5’-3’exonuclease in vivo, which is limited in extent to no more than 8-10 base pairs, and that repair is independent of RecA, RadA, and non-homologous end joining functions Ku and LigD. A mechanism of mycobacterial NucS MMR is discussed.

## Material and Methods

### Bacterial strains and media

The *M. smegmatis* strains used in this study were derived from mc^2^155; the *M. tuberculosis* strain used in this study was H37Rv. *M. smegmatis* and *M. tuberculosis* were grown in 7H9 broth with 0.05% Tween 80, 0.2% glycerol, and OADC (oleic acid-albumin-dextrose-catalase; Becton, Dickinson); transformants were selected on 7H10 plates with 0.5% glycerol and OADC with appropriate antibiotics. For the experiment shown in Fig. 3 involving the mismatch DNA repair specificity of NucS in vivo, *M. smegmatis* was transformed with pIR542. This plasmid is a Bxb1 *attP*-containing vector that expresses the Bxb1 phage Integrase and contains a defective *hyg* resistance gene. It was integrated into the endogenous Bxb1 *attB* site in the *M. smegmatis* chromosome by selection for zeocin resistance.

Deletion mutants of *M. smegmatis* were constructed by ORBIT (20) using *attB*-integrating plasmids pKM611 or pKM614, both of which contain a defective *hyg* gene (in opposite directions). *M. smegmatis* strains MGM199 (Δ*recA*) and MGM156 (Δ*Ku* and Δ*ligD*) were gifts kindly provided by Michael Glickman (21). For these strains, ORBIT was used to deliver pKM614 into the intergenic region between MSMEG_1397 and *rpsL*. In all cases, the defective *hyg* gene was placed in the chromosome in a direction that allowed oligos used in this study to target the lagging strand template. Verification of each ORBIT-promoted deletion strain was done by PCR amplification of both junctions of the inserted plasmid and the chromosomal target site, as well as the absence of an amplicon using primers internal to the target gene (compared to WT cells). Constructions of *leuB* mutants containing single or two base pair indels into *leuB* active site codon R216 were done using oligo-mediated recombineering as described previously (22). When needed, the following antibiotic concentrations were added to LB plates, 7H9 media or 7H10 plates: kanamycin (20 μg/ml), streptomycin (20 μg/ml), hygromycin (50 μg/ml), and zeocin (25 μg/ml). A list of mutants constructed and used in this study is shown in Table 1.

**Table 1:** Strains used and constructed for this study.

| Strain Name | Genotype | Description | Markers | Source |
| --- | --- | --- | --- | --- |
| MC<sup>2</sup>-155 | wild type | Efficient plasmid transformation (ept) | none | (37) |
| MGM199 | ΔrecA | Deletion of *recA* | none | (21) |
| MGM153 | ΔKu, ΔligD | Deletion of *Ku* and *ligD* | none | (21) |
| KM272 | Wild type<br>Defective Hyg<sup>R</sup> gene | pKM461 (RecT); pKM614 (defective *hyg* gene) inserted into the MSMEG_1397-*rpsL* intergenic region by ORBIT | Kan<sup>R</sup> Zeo<sup>R</sup> | This study |
| KM274 | ΔrecA<br>Defective Hyg<sup>R</sup> gene | MGM199/KM461 (RecT); pKM614 (defective *hyg* gene) inserted into the MSMEG-rpsL intergenic region by ORBIT | Kan<sup>R</sup> Zeo<sup>R</sup> | This study |
| KM295 | ΔKu, ΔligD<br>Defective Hyg<sup>R</sup> gene | MGM153/KM461 (RecT); pKM614 (defective *hyg* gene) inserted into the MSMEG-rpsL intergenic region by ORBIT | Kan<sup>R</sup> Zeo<sup>R</sup> | This study |
| KM165 | ΔnucS::pKM464 | MC<sup>2</sup>-155 deleted of MSMEG_4923 by Orbit | Hyg<sup>R</sup> | This study |
| KM183 | ΔnucS::attP | KM165 - cured of pKM464. | none | This study |
| VF20 | Wild type<br>Defective Hyg<sup>R</sup> gene | MC<sup>2</sup>-155/pKM461 (RecT); pIR542 (defective *hyg* gene) inserted into phage Giles site. | Kan<sup>R</sup> Zeo<sup>R</sup> | This study |
| VF22 | ΔnucS::attP<br>Defective Hyg<sup>R</sup> gene | KM183/pKM461 (RecT); pIR542 (defective *hyg* gene) inserted into phage Giles site. | Kan<sup>R</sup> Zeo<sup>R</sup> | This study |
| KM209 | ΔfenA::pKM464 | MC<sup>2</sup>-155 deleted of MSMEG_3883 by ORBIT | Hyg<sup>R</sup> | This study |
| KM210 | ΔfenA::attP | KM209 - cured of pKM464. | none | This study |
| KM211 | ΔfenA:pKM611<br>Defective Hyg<sup>R</sup> gene | KM210/pKM461 (RecT); pKM611 (defective *hyg* gene) Inserted at ΔfenA::attP site | Kan<sup>R</sup> Zeo<sup>R</sup> | This study |
| KM215 | $\Delta radA::pKM464$ | MC<sup>2</sup>-155 deleted of MSMEG_6079 by ORBIT | Hyg<sup>R</sup> | This study |
| KM284 | $\Delta radA::attP$ | KM215 - cured of pKM464. | none | |
| KM287 | $\Delta radA:pKM611$<br>Defective Hyg<sup>R</sup> gene | KM284/pKM461 (RecT); pKM611 (defective *hyg* gene)<br>Inserted at $\Delta radA::attP$ site | Kan<sup>R</sup> Zeo<sup>R</sup> | This study |
| KM287 | $\Delta recA, \Delta radA:pKM611$<br>Defective Hyg<sup>R</sup> gene | MGM199/pKM461 (RecT); pKM611 (defective *hyg* gene)<br>Inserted into $\Delta radA::attP$ site gene by ORBIT | Kan<sup>R</sup> Zeo<sup>R</sup> | This study |
| KM229 | MSMEG_2379<br>(leuB frameshift) | Two bp deletion in the arginine-101 codon; pKM461; pKM585 | Kan<sup>R</sup><br>Strep<sup>R</sup> | This study |
| KM230 | MSMEG_2379<br>(leuB frameshift) | One bp deletion in the arginine-101 codon; pKM461: pKM585 | Kan<sup>R</sup><br>Strep<sup>R</sup> | This study |
| KM231 | MSMEG_2379<br>(leuB frameshift) | One bp insertion in the arginine-101 codon; pKM461; pKM585 | Kan<sup>R</sup><br>Strep<sup>R</sup> | This study |
| KM234 | MSMEG_2379<br>(leuB frameshift) | One bp insertion in the arginine-101 codon; cured of ORBIT<br>plasmid pKM461 and pKM585. | none | This study |
| KM262 | NucS-D138A | Active site mutant of *M. smegmatis* NucS | none | This study |
| KM263 | NucS-E152A | Active site mutant of *M. smegmatis* NucS | none | This study |
| KM264 | NucS-K154A | Active site mutant of *M. smegmatis* NucS | none | This study |
| Mtb-25 | Mtb$\Delta nucS::pKM488$ | pKM461/H37Rv deleted of Rv1321 by ORBIT; cured of pKM461 | Hyg<sup>R</sup> | This study |
| Mtb-26 | Mtb $\Delta nucS::attP$ | Mtb-25 - cured of pKM488. | none | This study |

### Plasmids

Oligo-recombineering plasmid pKM402, Che9 RecT and Bxb1 Integrase-expressing plasmid pKM461, and Orbit integration plasmids have been described previously (20,23). Plasmid pIR540 is a Giles integration vector that expresses full-length *M. smegmatis nucS* gene (MSMEG_4923) from the PgroEL promoter. Plasmid pIR541 is similar to pIR540 except that it is missing the last five codons of the *nucS* gene.

Plasmid pIR542 is a Giles integration vector (Zeo^R^) that contains a defective hygromycin-resistant gene where the codon for glycine 110 (GGA) has been altered to stop codon (TAG). Plasmid pKM611 is a Bxb1 *attB*-containing ORBIT-integration vector that contains the same defective hygromycin-resistant gene from pIR542; pKM614 is the same as pKM611, but with the defective-hygromycin-resistant gene in the opposite direction relative to the Bxb1 *attB* site. Plasmid pKM585 contains an *attP* site, the defective *hyg* gene, and integrates into the endogenous *attB* site of *M. smegmatis* with the help of pKM461 (RecT, Integrase). Details of plasmid constructions, maps, and sequences are available upon request.

### Oligonucleotides

Oligonucleotides used for the determination of mismatch specificity are listed in Table S1. Oligonucleotides used for mutant construction by ORBIT are listed in Table S2 and were obtained from IDT as Ultramers at a concentration of 100 μM (delivered in 96-well plates). They were supplied desalted with no further purification and diluted 10-fold in sterile water. Final concentrations (250 to 350ng/ml) were determined by absorbance maxima at 260 nm (Abs_260_). Oligos used in RecT-promoted recombineering experiments for mutant constructions and repair of the defective *hyg* gene were obtained from Life Science Technologies and are listed in Table S2; 1-2 ug of each oligo was typically added to electrocompetent cells for SNP transfer.

### Transformations

Details of the transformation procedure using the ORBIT technology for mutant strain construction have been described previously (20). Electroporations for oligonucleotide-promoted recombineering experiments, here to restore the defective *hyg* gene, have been described previously (22). Briefly, 150 ul of fresh overnight culture of *M. smegmatis* was added to 20 ml of 7H9-OADC-tween containing 20 μg/ml kanamycin and swirled at 37°C overnight. The following day, when the culture reached an absorbance of 0.5, anhydrotetracycline (Atc) was added to a final concentration of 500 ng/ml. In cases where the culture went past an absorbance of 0.5, it was diluted back to 0.5 and Atc was added. The culture was allowed to grow for an additional three hours or until the O.D. reached 1.0. Cells were then collected by centrifugation, swirled on ice for 10 minutes, and washed twice with 20 ml of cold 10% glycerol. After the final centrifugation, the cells were resuspended in 2 ml 10% glycerol and kept on ice. Oligonucleotides (2 μg) were placed in Eppendorf tubes and 380 μL of electrocompetent cells were mixed with oligos then transferred to 0.2 cm electroporation cuvettes. Cells were shocked and transferred to 2 ml 7H9-OADC-tween and grown overnight at 37°C as described previously (20). Appropriate dilutions of overnight cultures were plated on LB plates or 7H10-OADC plates (+/-50 ug/ml hygromycin) to determine the fraction of Hyg^R^ transformants.

### Mutagenicity Assay

The lack of NucS repair activity in M. *smegmatis* was determined by plating 200 μL of a saturated culture (O.D. of 1.0-1.5) on 7H10 plates containing 150 ug/ml of rifampicin; total cell numbers were determined by plating on 7H10 plates. Plates were incubated for 4-5 days at 37°C. The frequency of spontaneous mutation was determined by the titer of rifampicin-resistant colonies divided by the total cell titer.

### Recombineering with oligonucleotides generating two mismatches

To examine the extent of resection after NucS cutting, we designed nine oligos to create G-G and C-T mismatches separated by 4 to 32 base pairs along the chromosome, allowing us to measure the extent of processing of the C-T mismatch as a function of the distance (and position) from the NucS-proposed dsDNA cut (or nick) at the G-G mismatch. The sequence of these oligos and the position of the G-G mismatch relative to the C-T mismatch in each oligo (following annealing to the lagging strand template) is shown in Fig. 7A. These oligos were electroporated into *M. smegmatis* containing pKM461 following induction with anyhydrotetracycline, as described above. After overnight growth, portions of the culture were plated on 7H10 and 7H10 plates with 50 μg/ml hygromycin to determine the frequency of Hyg^R^ transformants.

## Results

### Mutator phenotype of *M. smegmatis nucS* strain and testing of the PIP domain

Using the previously described ORBIT method for the generation of chromosomal modifications in mycobacteria (see Materials and Methods), a deletion mutant of *M. smegmatis nucS* gene (MSMEG_4923) was constructed. The frequency of spontaneous mutation to rifampicin resistance was measured in both wild type *M. smegmatis* and its ΔnucS derivative (see Fig. 1). As reported previously (19), an *M. smegmatis* strain deleted of *nucS* exhibited a mutator phenotype, with a greater than 100-fold increase in the frequency of spontaneous mutations when compared to a wild type strain.

**Figure 1:**
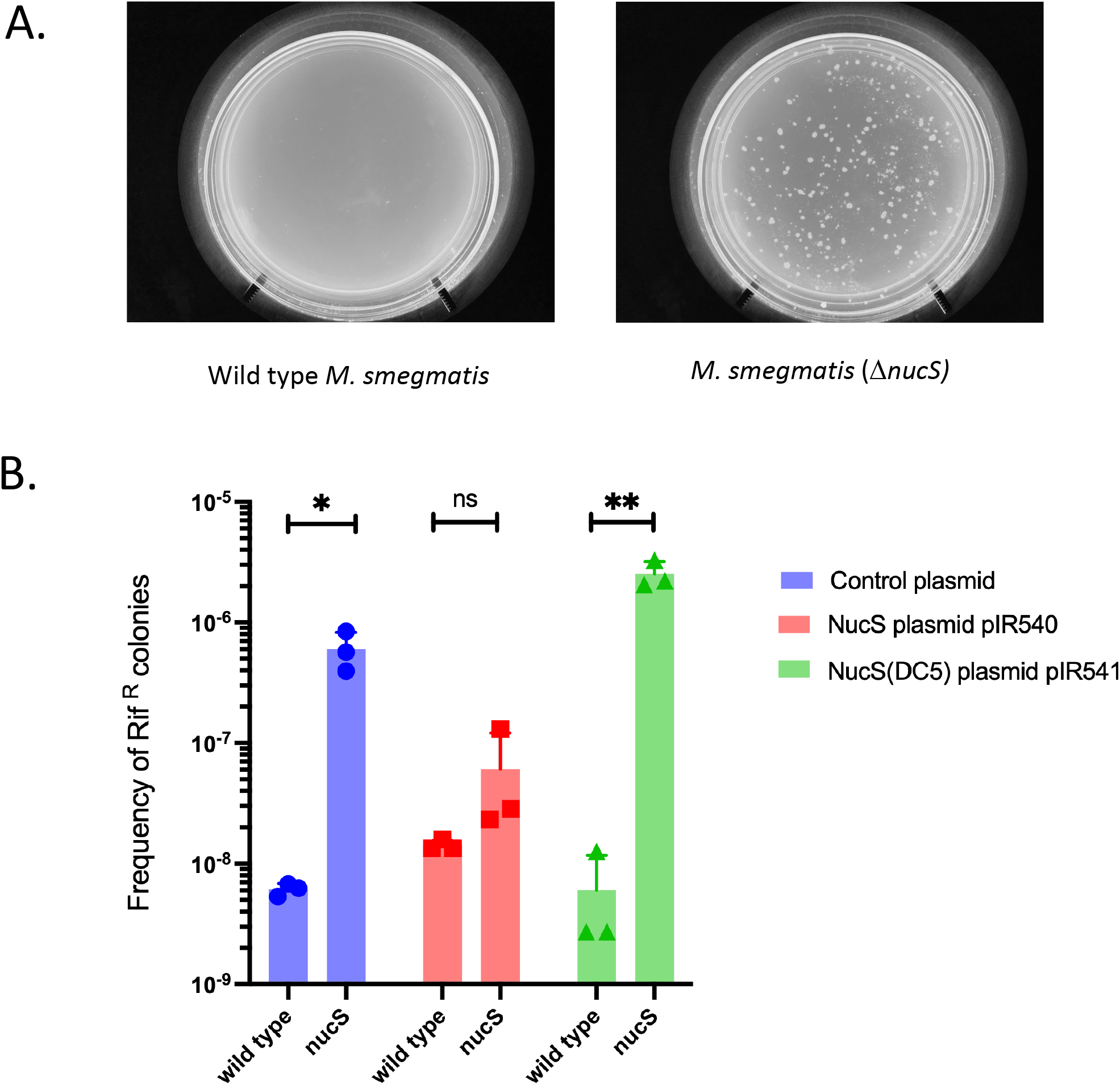
Mutagenic phenotype of *M. smegmatis* Δ*nucS* strain. **(A)** Wild and a Δ*nucS* derivative of *M. smegmatis* were grown in 7H9-OADC-tween to saturation (O.D. of 1.5) and 200 ul of cultures were spread on 7H10-OADC-tween plates containing 150 ug/ml rifampicin. **(B)** *M. smegmatis* wild type and ΔnucS strains containing a control plasmid, pIR540 (expressing full-length NucS), or pIR541 (expressing NucS containing a 5 amino acid truncation from the C-terminus – NucS(DC5) were grown to saturation and plated on 7H10-OADC-tween rifampicin plates as described in above. The mutation frequency is the titer of rifampicin-resistant mutants divided by the total number of cells plated. Data represent the means ± SD from three biological replicates; two-tailed *t*-test was performed (\**P* < 0.05).

In complementation experiments, the expression of NucS from pIR540 reduced the mutation frequency considerably (about ∼12-fold), close to that observed in WT cells. However, a similar construct expressing a NucS protein with the last five amino acids deleted from the C-terminus (NucS-ΔC5) failed to complement the *nucS* strain, resulting in high levels of mutagenesis (see Fig. 1B). This region of *M. smegmatis* NucS is homologous to the C-terminal region in *P. abyssi* NucS that has been shown to interact with the PCNA sliding clamp, thus called the PIP domain (for <u>P</u>CNA interacting <u>p</u>rotein) (12). This region is also identical to the C-terminal 5 amino acids of NucS from *C. glutamicum*, which required an interaction with the β-sliding clamp to promote dsDNA cutting of mismatched-containing substrates in vitro (17,18). The failure of the *M. smegmatis* NucSΔC5 to suppress the mutagenic phenotype of the ΔnucS mutant suggests that an interaction of the C-terminal PIP domain in *M. smegmatis* NucS with the β-sliding clamp at the replication fork is required for efficient NucS-promoted cutting in vivo, and thus subsequent MMR.

It is possible that the loss of the C-terminal 5 amino acids causes *M. smegmatis* NucS protein to become unstable resulting in a mutagenic phenotype. To address this issue, we performed ORBIT to tag *nucS* to be able to follow the protein and its truncated version by Western analysis. However, C-terminal chromosomal fusions of wild type *nucS* with either Flag-DAS or GFP tags resulted in strains with mutagenic phenotypes (data not shown), presumably by interfering with the interaction of the NucS PIP domain and the β-clamp. Tagging of chromosomal *nucS* at the N-terminus was then performed using a 6x-His tag, but this modification also resulted in a strain that exhibited a mutagenic phenotype, suggesting the His tag may have interfered with dimerization of NucS or the binding of a protein partner in vivo. These results indicate that an interaction between a free-ended C-terminus PIP domain of *M. smegmatis* NucS and the β-clamp is required for efficient mismatch repair in vivo, and likely explains the inability of purified *M. smegmatis* NucS alone to recognize and cut mismatched dsDNA substrates in vitro, as was observed in a previous study (19). Biochemical analysis of NucS with mycobacterial DnaN β-clamp and mismatched substrates will be needed to verify this supposition.

### Mismatch DNA repair specificity in vivo

Previous mutational accumulation studies in both *C. glutamicum* and *M. smegmatis* have shown that the absence of the NucS MMR system results in an a greatly elevated rate of transitions, both G:C > A:T and A:T > G:C (the latter one being the higher of the two, opposite to what is observed in WT cells). In particular, Castañeda-Garcia *et al* (24) used an integrated plasmid reporter system to measure the mutational rates (and specificities) of both transitions and transversions using mutant versions of *aph* (kan^R^) alleles. With their reporter strains, the fold increase in mutation rates (ΔnucS/WT) was highest (65-fold) for transitions that required repair of a G-T mismatch to generate a Kan^R^ cell. In contrast, the fold increase in transversion mutations on their reporter plasmids was low, including ones that are predicted to generate G-G and T-T mismatches (2.4 and 3.6-fold respectively).

However, these mismatches are recognized and cut efficiently by *C. glutamicum* NucS in vitro. The low rate of NucS-dependent activity on transversion repair, despite efficient cutting of G-G and T-T mismatches in vitro, may be simply due to the low levels of G-G and T-T mismatches that arise during DNA replication in mycobacteria. Or it may mean that NucS does not act on these mismatches at high efficiency in vivo, despite highly efficient cutting of them in vitro.

We were thus encouraged to set up a similar assay to examine the mycobacterial mismatch repair system in vivo to compare the specificity of mismatch recognition to what has been reported in vitro. We thus tested the ability of Che9 RecT-mediated oligonucleotide recombineering to generate defined mismatches in a way that would allow us to follow the fate of mismatches by simple plating assays. A first test of this system was to deliver mismatches to the replication fork within the *rpsL* gene (22,25). Oligos were designed to generate either K43R or K43N changes to the *rpsL* ribosomal protein, both of which are known to confer resistance to streptomycin (26). The K43R oligo creates a C-A mismatch at the replication fork, while the K43N oligo generate at G-G mismatch (see Fig. 2). According to mismatch specificity reported for *C. glutamicum* NucS in vitro, the K43R oligo would create a mismatch in *M. smegmatis* that would not be subject to NucS-mediated repair (C-A) and thus generate high levels of streptomycin-resistant colonies. Along the same lines, the K43N oligo would generate a mismatch that is subject to repair (G-G) and thus fail to generate streptomycin-resistant colonies. These predictions were fulfilled by the data shown in Table 2, showing that recombineering can be used to deliver mismatches to test in vivo specificities, and that *M. smegmatis* shows a similar mismatch repair specificity as exhibited in vitro by *Actinobacteria*, but do so in vivo. The same experiment was performed in *M. tuberculosis* with similar results (Fig. S2).

**Figure 2:**
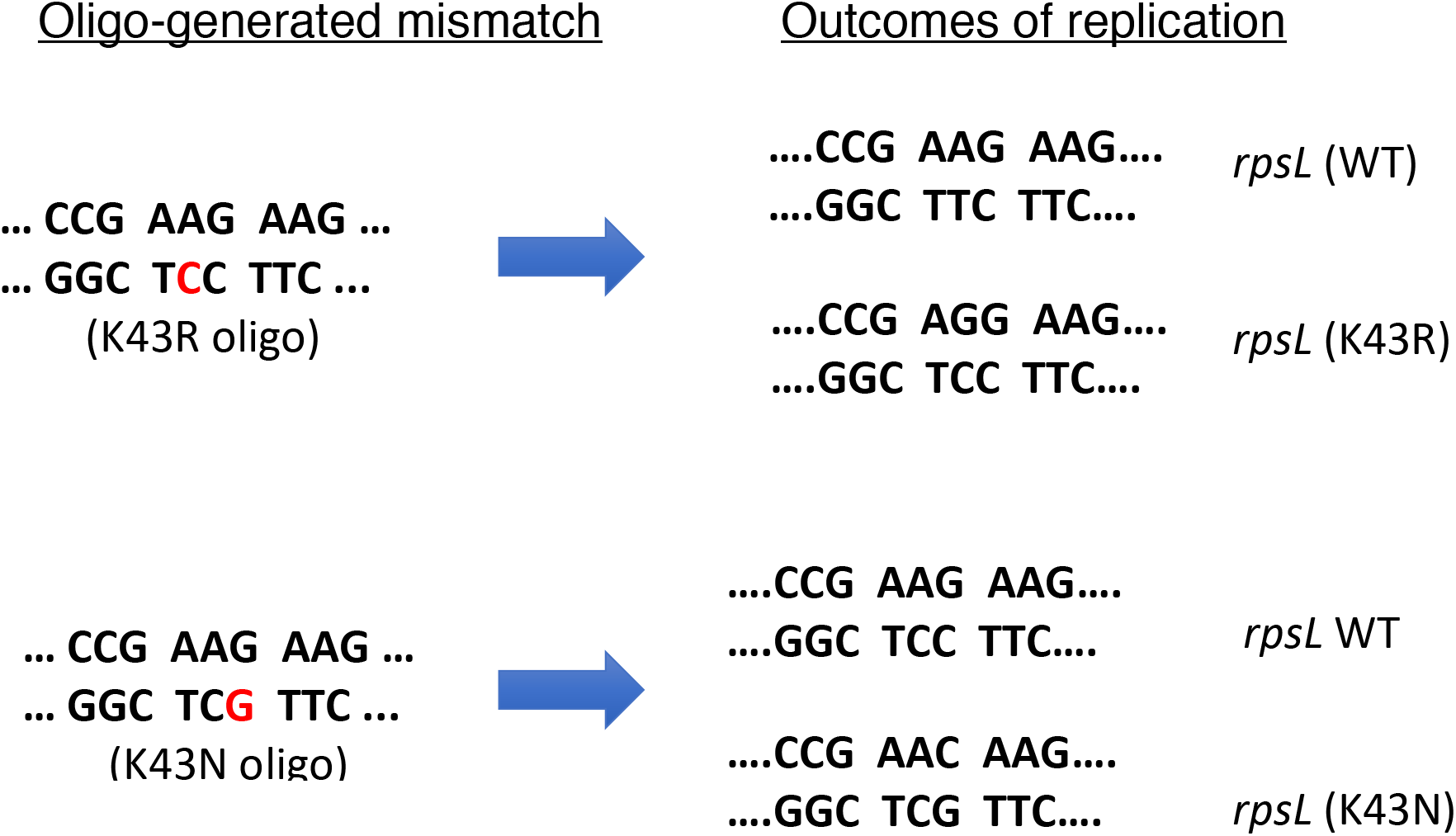
Oligo-mediated recombineering of the *rpsL* locus of *M. smegmatis*. (Left) Sequences of the oligo-generated mismatch in the *rpsL* gene resulting in a mismatch (A-C) predicted not to be recognized by mycobacterial NucS, and a mismatch (G-G) expected to be acted upon by NucS, as predicted from in vitro results of *C. glutamicum* (*17,18*). (Right) Following replication of mismatched-containing DNA, both wild type and mutant versions of *rpsL* are generated. Both mutants are known to cause streptomycin resistance in mycobacteria (26).

**Table 2:** Frequencies* of oligonucleotide-mediated streptomycin resistance in *M. smegmatis*.

| Strain | K43R oligo<br>(A/C mismatch) | K43Q oligo<br>(G/G mismatch) |
| --- | --- | --- |
| Wild type | 1.59 | 0.0097 |
| $\Delta nucS$ | 1.05 | 1.4 |

\* Titer of streptomycin resistant colonies/total colonies ( $\times 10^{-4}$ )

To further test the in vivo specificities of mismatch repair, a second assay was designed to study all the other types of mismatches. For this purpose, we inserted a stop codon (TAG) in place of the glycine codon (GGA) at position 110 in the hygromycinB phosphotransferase gene (*hyg*) in plasmid pIR542, a Giles integrating plasmid. The G110 residue was selected because of its presence in an extended loop on the surface of the protein (see Fig. S1A). In addition, the G-110 residue is not conserved among *hyg* gene from other bacterial species (Fig. S1B), highlighting this position as one that is likely tolerant to multiple types of amino acid substitutions. This prediction was fulfilled in the course of these assays.

Oligos were then designed to convert the stop codon in the integrated plasmid pIR542 to various sense codons, allowing for full translation of the *hyg* gene. Each oligo generates a different type of mismatch (see Table S1). If a mismatch escapes repair, it leads to a recombinant that expresses a functional phosphotransferase and generates Hyg^R^ colonies; on the contrary, if the mismatch is repaired, no or few Hyg^R^ transformants would be observed.

*M. smegmatis* containing pKM402 (RecT producer) and pIR542 were electroporated with the oligos listed in Table S1 and the outgrowths were plated on LB plates with and without hygromycin, as described in Materials and Methods; the results are shown in Fig 3. In otherwise wild type *M. smegmatis,* oligos that generated G-G, G-T and T-T mismatches generated Hyg^R^ colonies more than 500-fold lower frequency relative to transformations with oligos that generated all the other types of mismatches, indicative of repair of G-G, G-T and T-T mismatches in vivo. All the other mismatches tested escaped MMR, generating Hyg^R^ colonies at frequencies 10,000-fold greater than the control transformation with no oligo (Fig. 3A). When the same experimental protocol was performed in the *M. smegmatis* Δ*nucS* strain, transformation with all the oligos generated high frequencies of Hyg^R^ colonies, revealing that in the absence of MMR, all mismatches generated were incorporated into the chromosome (Fig. 3B). These results reveal that *M. smegmatis* shows the same mismatch repair specificity in vivo as the archaeal TKO and the actinobacterial *C. glutamicum* strain exhibited in vitro, indicating that these MMR systems recognize identical types of mismatches and likely behave by similar mechanisms. Additionally, the G-T mismatch recognition specificity revealed here is consistent with the mutational specificity of *M. smegmatis* reported by Castaneda-Garcia *et al*. (24), who showed a high rate of transitions when the reporter plasmids generated G-T mismatches in order to restore kanamycin resistance. On the other hand, G-G and T-T (which if unrepaired lead to transversions), do not show high levels of NucS-dependent repair in mutational accumulation studies, even though NucS recognized and repaired G-G and T-T mismatches in vivo at the same rate as G-T mismatches (Fig. 3A), This result is further elaborated on in the Discussion section.

**Figure 3:**
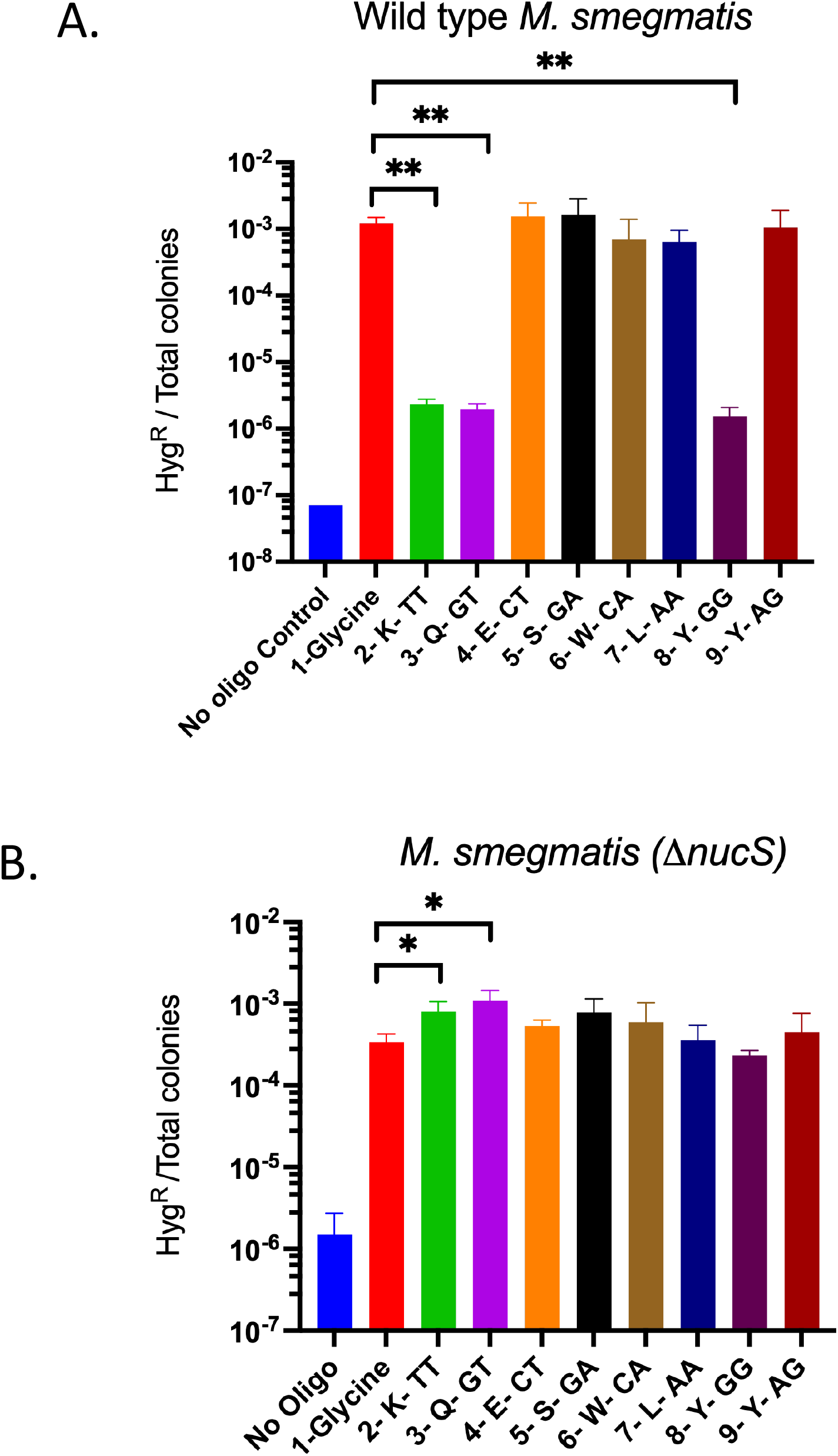
Mismatch recognition specificity of *M. smegmatis* NucS in vivo. Oligo-mediated recombineering experiments were carried out in WT **(A)** or Δ*nucS* **(B)** *M. smegmatis* cells containing RecT-producer pKM402 and a defective hygromycin gene, where codon-110 for glycine (GGA) had been altered to a stop codon (TGA). The oligos used generated 7 different mismatches for restoration of the *hyg* gene. Repair and non-repair of the mismatches are evident by the appearance of low and high frequencies of hygromycin-resistant colonies, respectively. Experiments were done in triplicate. Data represent the means ± SD from three biological replicates; two-tailed *t*-test was performed comparing results from the oligo that restores the original glycine codon to position 110 (red bar) to all other oligos that created the defined mismatches; \**P* < 0.05. Only significant *p* values are shown.

Inability of *M. smegmatis* NucS to repair small indels.

A test to examine the in vivo repair capacity of *M. smegmatis* NucS on small indels was done with our recombineering assay, examining the ability to remove 1-2 bp insertions and deletions in the *leuB* gene. Such indels inactivate *leuB* activity and recombineering with oligos that removed these indels allow this strain to grow on 7H10 plates without leucine supplementation. If NucS is inactive on DNA containing 1-2 bp indels, such oligo-mediated leuB^+^ strains will be easily observed. This was, in fact, the case, as insertions and deletions between 1-2 bp were easily removed from these *leuB* frameshift mutants using an oligos that contained the wild type *leuB* sequence, restoring the defective *leuB* allele back to wild type at high frequencies (see Fig. 4). The lack of activity of mycobacterial NucS on indels in this recombineering assay is in agreement with the MA experiments and reporter plasmids previously described (24). This result, and those in Fig 3, further reveal that delivery of DNA mismatch to the chromosome by oligo-mediated recombineering faithfully mimics *bona fide* DNA mismatches that occur in the replication fork in vivo.

**Figure 4:**
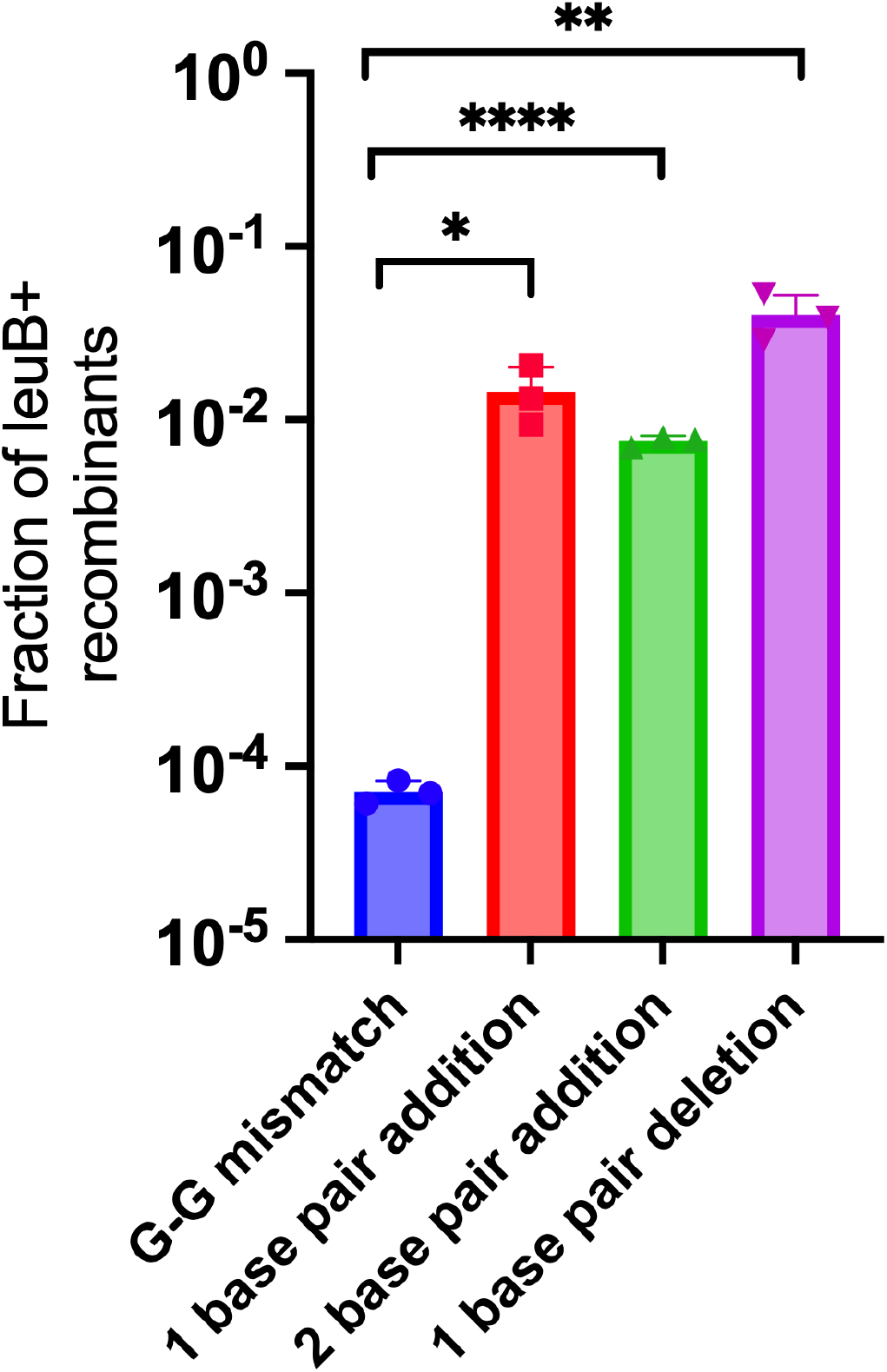
NucS does not repair indels. Oligo-mediated recombineering was performed in strains where the *leuB* gene had been deactivated by the deletion of 1 or 2 base pairs, or by the insertion of 1 base pair in the codon that encodes for the active site residue Arg-216. An oligo containing the wild type *leuB* sequence was electroporated into these *leuB* cells which expresses the RecT annealase from plasmid pKM461. LeuB+ recombinants were measured by plating the cells on 7H10-OADC-tween plates without leucine supplementation; total cells were titered on 7H10-OADC-tween plates with 40 ug/ml leucine. For comparison, the result for a *leuB* strain is shown where the active site codon was disabled and could be efficiently restored to wild type with an oligo that creates a repairable G-G mismatch in Δ*nucS* host, but not in wild type cells. Data represent the means ± SD from three biological replicates: Student’s two-tailed *t*-test, \**P* < 0.05.

### Identification of likely NucS nucleolytic active site residues

Nakae *et al* (14) solved the structure of the NucS protein from *P. abyssi* and identified three amino acid residues that may be involved in nucleolytic activity (Fig. 5A). In comparison with the *P. abyssi* NucS protein sequence, the corresponding amino acids in *M smegmatis* NucS are D138, E152 and K154. Using Che9 RecT-mediated oligo recombineering, each of these residues was separately changed to alanine and the subsequent strains were tested for mutagenic activity by plating on 7H10 plates containing rifampicin. The D138A, E152A, and K154A mutations each conferred a mutagenic phenotype to *M. smegmatis*, implicating all these residues in NucS endonucleolytic activity (Fig. 5B). Further confirmation of these residues as nucleolytic active site residues will require biochemical analysis of the mutant proteins.

**Figure 5:**
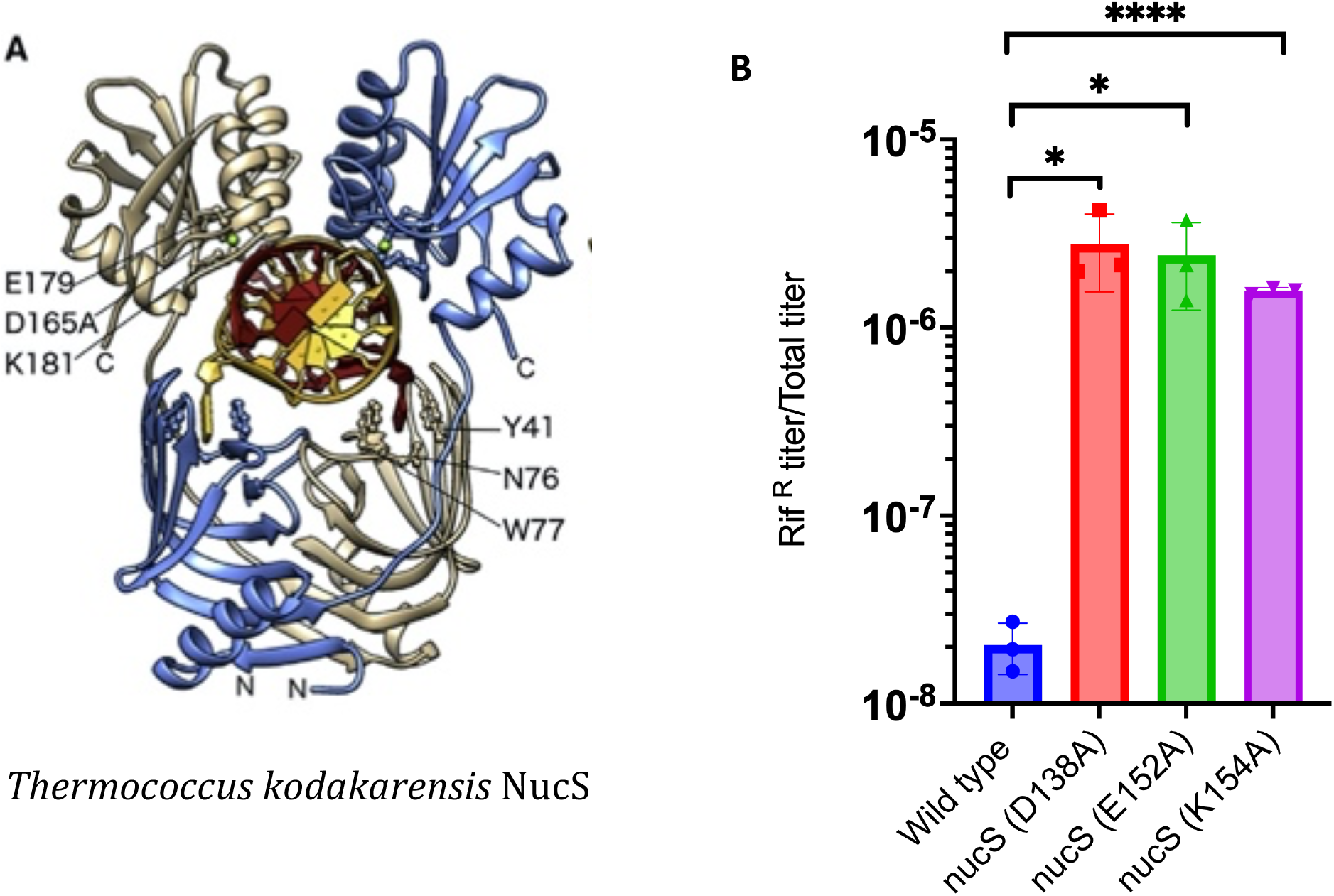
Mutations suspected of being critical for exonucleolytic activity in NucS are defective for MMR. (A) The structure of the NucS protein from *Thermococcus kodakarensis* bound to dsDNA highlighting residues thought to be critical for exonucleolytic activity: E179, D165, and K181. (B) The corresponding residues in *M. smegmatis* NucS were changed to alanine by recombineering and tested for levels of spontaneous mutation to rifampicin. Data represent the means ± SD from three biological replicates: Student’s two-tailed *t*-test, \**P* < 0.05. The *T. kodakarensis* NucS structure was reprinted from Nakae et. al. (14) with permission.

### Resection of NucS-promoted cuts in vivo involves limited processing by a 5’-3’ exonuclease

The identification of nucleolytic active site residues in *M. smegmatis* NucS that lead to high mutability suggests that cutting in vivo likely occurs in the processing of mismatches in mycobacteria. Whether this involves a ssDNA nick (as seen with the MutSLH system) or a dsDNA break (as suggested by in vitro results from *C. glutamicum* NucS_Cg_) has not currently been established in vivo. In either case, we sought to examine the processing of such cuts by endogenous exonucleases in vivo by delivering two mismatches (within one oligo) to the chromosome via recombineering, with one mismatch (C-T) generating Hyg^R^ mutants (as described above) and the other mismatch (G-T) being recognized and cut by NucS. The oligo sequence is shown in blue in Fig. 6B and generates mismatches that are 32 bp apart on the lagging strand. It is assumed that exonucleolytic processing would occur from the NucS-induced cut at codon 99 to codon 110 of the Hyg^R^ gene. If processing by a 5’-3’exonuclease proceeds past codon 110 of the Hyg^R^ gene (Fig. 6C), subsequent repair by resynthesis of the gap (or by recombinational repair) would lead to loss of the C-T mismatch and restoration of the stop codon. This, in turn, would lead to loss of Hyg^R^ transformants relative to the use of an oligo that did not generate a repairable mismatch (control oligo), but still generates the C-T mismatch that repairs the defective hyg gene. On the contrary, limited exonucleolytic processing from the G-T mismatch at codon 99 that did not proceed to codon 110 would not affect the level of Hyg^R^ transformants compared to the control oligo (Fig. 6D).

**Figure 6:**
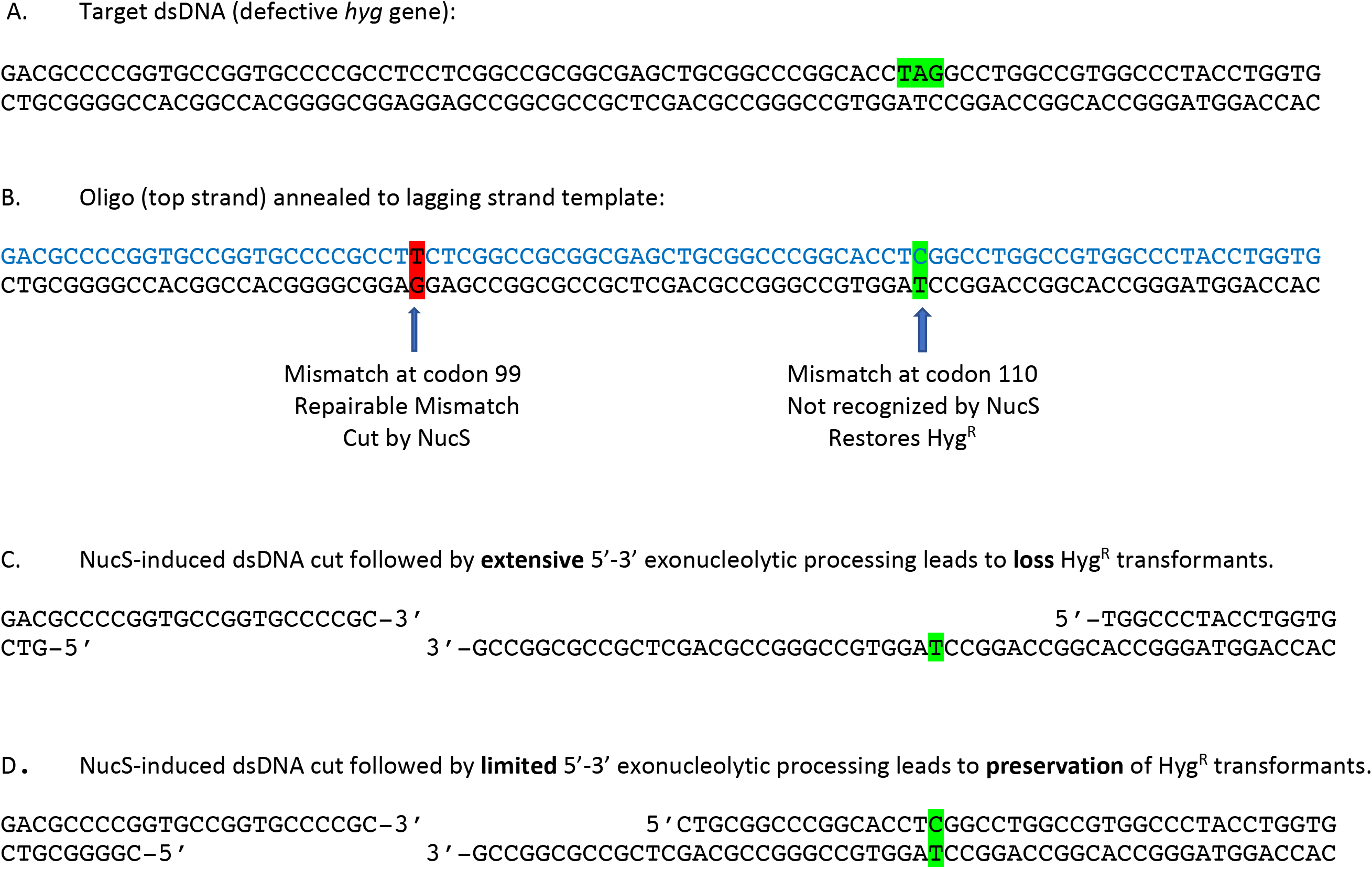
Description of a double mismatch-generating oligo. (A) The dsDNA sequence of the defective hygromycin-resistant gene (*hyg*). The GGA-110 codon has been altered to stop codon TAG (highlighted in green). (B). After annealing of the oligo (in blue type) to the target site in the *M. smegmatis* chromosome, the oligo creates two mismatches: a C-T mismatch (unresponsive to NucS) that leads to high levels of Hyg^R^ colonies and a T-G mismatch that is acted upon by NucS in vivo. (C) NucS is predicted to make a dsDNA cut at the mismatch. If the ends of the breaks are acted upon by a 5’-3’ exonuclease, Hyg^R^ recombinants will be lost if the processing extends to the C-T mismatch or beyond and removes the “C”. (D) If the processing by a 5’-3’ exonuclease is limited and does not extend to the C-T mismatch, high levels HygR transformants will be generated.

A first test of this two-mismatch-promoting oligo design showed that it could generate Hyg^R^ transformants at the same frequency as a control oligo that generated the C-T mismatch but not the G-G mismatch (∼ 5 x 10^-4^ Hyg^R^ transformants/total cell titer – data not shown). Sequencing of PCR fragments containing this region from 8 Hyg^R^ colonies found that all the G-T mismatches were repaired (resulting in a “C” in the replicating strand). This result shows that exonucleolytic processing from the NucS-promoted cut at the G-T mismatch did not extend 32 bases to the C-T mismatch, leaving the frequency of Hyg^R^ colony formation intact relative to the control oligo (no G-G mismatch).

Given the result above, we performed the same test with a series of oligos that varied the distance between a repairable mismatch (this time G-G) and the C-T mismatch that confers Hyg^R^. Oligos containing both a G-G and C-T mismatch were electroporated into *M. smegmatis* containing pKM461 (RecT) and the defective *hyg* gene; see Fig. 7A for positions of the G-G mismatches relative to the C-T mismatch in each of these oligos. We followed the frequency of Hyg^R^ colonies from the outgrowth when the two mismatches were relatively close (4-5 bp) or further apart (16 or 20 bp), as well as oligos where the repairable mismatch was downstream of the C-T mismatch at variable distances (see Fig.7A). At a distance 20 bp upstream of codon 110, a G-G mismatch-guided NucS-promoted cut did not lead to loss of Hyg^R^ colonies relative to the oligo with no G-G mismatch. This result suggests that 5’-3’ processing of the predicted dsDNA cut in the top strand did not extend 20 bp to the C-T mismatch, which would have resulted in loss of Hyg^R^ colonies relative to the oligo with no G-G mismatch (similar to the result described above). On the other hand, when the G-G mismatch was only 5 bp away from the C-T mismatch, there was a 50-fold drop the frequency of Hyg^R^ colonies. This result suggests that a dsDNA break at the G-G site close to the C-T mismatch resulted in 5’-3’ exonucleolytic processing of the top strand that extended to the C-T mismatch, removing the “C” base, with eventual restoration of an “A” at this position to restore the stop codon (leading to loss of Hyg^R^ colonies) (see Fig 7B).

**Figure 7:**
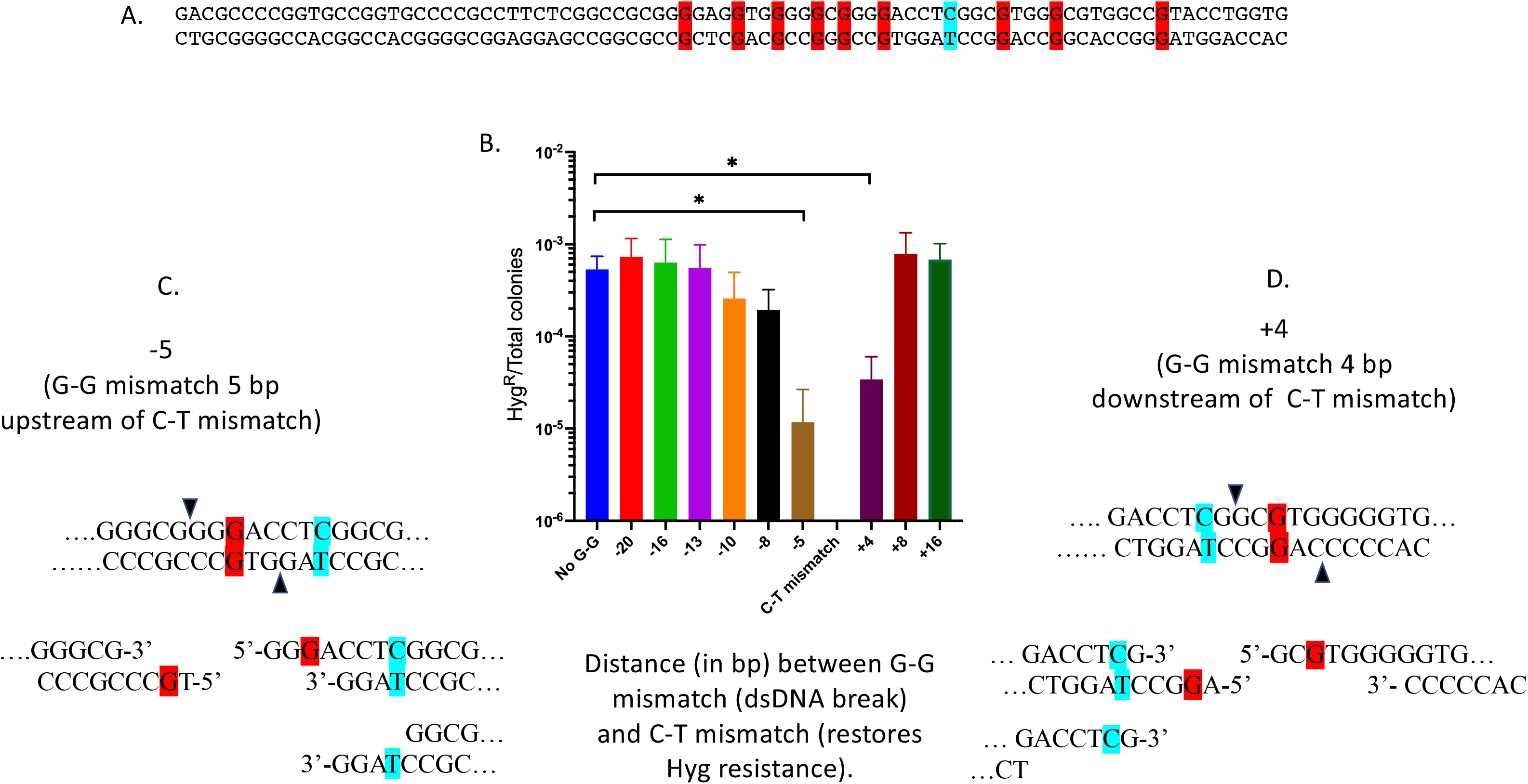
5’-3’ exonuclease processing of NucS-induced breaks. **(A)** The target region of the defective *hyg* gene is shown; the C-T mismatch leading to Hyg^R^ is shown in cyan. The top strand is the sequence of the recombineering oligo (replicating lagging strand) and the positions where nine different oligos produce a repairable G-G mismatch are shown in red. **(B)** Oligo recombineering was carried out in *M. smegmatis* carrying the defective *hyg* gene and plasmid pKM461(RecT). Each of nine oligos used in this assay generates a C-T mismatch at a fixed position (*i.e.,* the stop codon) and a G-G mismatch at variable positions as described in panel A. The names of the oligos on the X-axis define the number of base pairs between the two mismatches, and if the G-G mismatch is upstream (-) or downstream (+) of the C-T mismatch. The frequency of Hyg^R^ colonies and total cell number were determined by plating on 7H10-OADC-tween plate with and without hygromycin, respectively. Data represent the means ± SD from three biological replicates; two-tailed *t*-test was performed comparing results from the control oligo (no G-G mismatch, blue bar) to each test oligo; \**P* < 0.05. Only significant *p* values are shown. **(C)** Diagram of a dsDNA cut made by oligo (−5) *i.e.*, 5 base pairs upstream of the C-T mismatch that conveys Hyg^R^. **(D)** Diagram of a dsDNA cut made by oligo (+4) *i.e.*, 4 base pairs downstream of the C-T mismatch that conveys Hyg^R^.

When the position of the G-G mismatch was designed to be downstream of the C-T mismatches (three positions shown on Fig. 7A), the assay can follow the exonucleolytic processing of the 5’ overhang in the bottom strand (Fig 7B and 7D). In this case, exonucleolytic processing of the bottom strand (toward the C-T mismatch) would remove the “T” base, leaving the “C” in the top strand, securing the replacement of the stop codon with a TCG codon (Arg), leading to high levels of Hyg^R^. Thus, in this configuration, one would expect no (or less) loss of Hyg^R^ colonies with any oligo containing a G-G mismatch downstream of the C-T mismatch. This outcome is what was observed for oligos 8 and 16 bp downstream of the C-T mismatch, and partially true when the G-G mismatch was positioned only 4 bp downstream from the C-T mismatch (see Fig. 7B and 7D). In this latter case, there was a ∼20-fold drop in Hyg^R^ colonies, instead of the 50-fold drop seen when the G-G mismatch was located on the other side of the C-T mismatch) The key observation from the above assay is that processing of the break induced by NucS is principally performed by a 5’-3’ exonuclease. This can be seen with the 3-fold higher loss of Hyg^R^ colonies with G-G at the −5 position (relative to C-T) compared to the +4 position. If processing were performed by 3’-5’ exonucleases, the opposite would have been expected to be observed. Furthermore, given the 5’ overhangs predicted from a NucS-promoted dsDNA cut made at the G-G mismatch at the +4 position, nibbling of only 2 bp of the 3’ end would result in loss of Hyg^R^ transformants (see Fig. 7D). However, processing of the G-G mismatch at the −5 position requires at least 8 base pairs to be excised before loss of any Hyg^R^ colonies is observed (see Fig. 7C). Nonetheless, Hyg^R^ colony loss occurs at a 3-fold higher frequency at the −5 position relative to the +4 position.

These results support a model of 5’-3’ exonucleolytic processing of the dsDNA break at the site of a NucS-induced cut at a mismatch in vivo. What is also revealing is that while the data show a gradual increase in Hyg^R^ colonies as the G-G mismatch is moved from the −5 to the −13 position (though not statistically significant), at −20 bases away the frequency of Hyg^R^ transformants is equal to the frequency seen with the control oligo (*i.e*., no G-G mismatch). This result suggests that while there is 5’-3’ exonucleolytic processing from the NucS-induced cut, it is very limited to a small patch of DNA and inconsistent with the at least ∼200 bp to 1 kb of processing of ssDNA that usually accompanies RecA-promoted dsDNA break repair events. Such short patch repair is reminiscent of a similar pathway of mismatch repair events promoted by MutS in *E. coli*. While these results have been presented in Figure 7 in the context of NucS making a dsDNA cut at the G-G mismatch, the same conclusions can be drawn if NucS promoted a nick in the replicative strand (blue strand in Fig. 6B).

### Repair of NucS-promoted ds-DNA breaks is not dependent on RecA-or RadA-promoted homologous recombination, or NHEJ

We sought to use recombineering to deliver a repairable mismatch (G-G) in different mutant strains, so that we could test directly for genetic functions that work downstream of NucS (*i.e.,* ones that would process a dsDNA break, for example *recA*). However, simply targeting the defective *hyg* gene (as was done in Fig. 3) could not be used in such an assay, as the repair and non-repair of a G-G mismatch (*after* NucS cutting) gives the same result, *i.e.,* loss of Hyg^R^ colonies (relative to a control oligo containing a non-repairable mismatch).

Thus, we turned to the double-mismatch-promoting oligo assay described above to examine the role of homologous recombination in NucS-promoted MMR. If a dsDNA cut is indeed generated in vivo at the G-G mismatch, and recombination repair acts on that break, the absence of RecA protein would be expected to lead to an unrepaired dsDNA break and death of that cell, resulting in loss of Hyg^R^ transformants relative to a WT host. On the other hand, efficient repair of the G-G mismatch in this assay has no effect on the levels of Hyg^R^ colonies since 1) it is far enough away (>20 bp) to not interfere with the C-T mismatch (see Fig 7), and 2) the repair event is in the wobble position of the targeted codon so as not to affect the amino acid residue at that position. Thus, loss of Hyg^R^ transformants in a mutant strain, compared to a control oligo not generating the G-G mismatch, would indicate involvement of that gene in MMR.

Besides RecA, the *M. smegmatis* RadA function has also been considered to be a candidate for repair of an MMR-induced dsDNA break. RadA is an archaeal analog of the RecA recombinase and can form nucleoprotein filaments on DNA and catalyze DNA pairing and strand exchange (27,28). RadA has a role in stimulating branch migration that extends heteroduplex formations in RecA-mediated strand transfer reactions (29,30). Mutants of RadA in *E. coli* have strong synergistic phenotypes with *recG* mutants, a branch migration enzyme, and show synthetic genetic interactions for survival to AZT when combined with several known helicases including PriA, RuvAB, UvrD, and others (31). *B. subtilis* RadA has also been shown to possess a 5’-3’ helicase activity capable of unwinding recombinational intermediates (32). The *M. smegmatis radA* gene seemed especially relevant to a study of *nucS* function since both these genes are of archaeal origin and in one genus of archaea (*Thermococcus*), *nucS* is co-transcribed with *radA* (12,13).

We thus tested both recA, radA, and the double mutant in the double-mismatch recombineering assay; the results are shown in Fig. 8. In support of the data of limited resection shown in Fig. 7, neither *recA*, *radA*, or the double mutant exhibited a loss in the frequency of Hyg^R^ transformants following transfer of the oligo generating the G-G mismatch relative to the control oligo (where a G-G mismatch is not generated). In *recA* hosts, there was actually a small stimulation of the frequency of Hyg^R^ cells. It is not known why this is the case, as it could be due to an inhibitory effect of RecA on NucS-promoted repair or may be the result of increased RecT-promoted annealing of the oligo in the absence of RecA. Nonetheless, since no loss of Hyg^R^ transformants is observed with these mutants, recombinational repair is not involved in NucS-promoted MMR. Further proof that MMR occurred in the absence of both RecA and RadA came from sequencing of the *hyg* gene from eight Hyg^R^ transformants (4 from *recA* cells and 4 from the *recA radA* double mutant), all of which showed that the G-G mismatch had been efficiently repaired, while in 4 *nucS* control cells, the “G” for “C” SNP was efficiently transferred (*i.e.,* the mismatch had escaped MMR).

**Figure 8:**
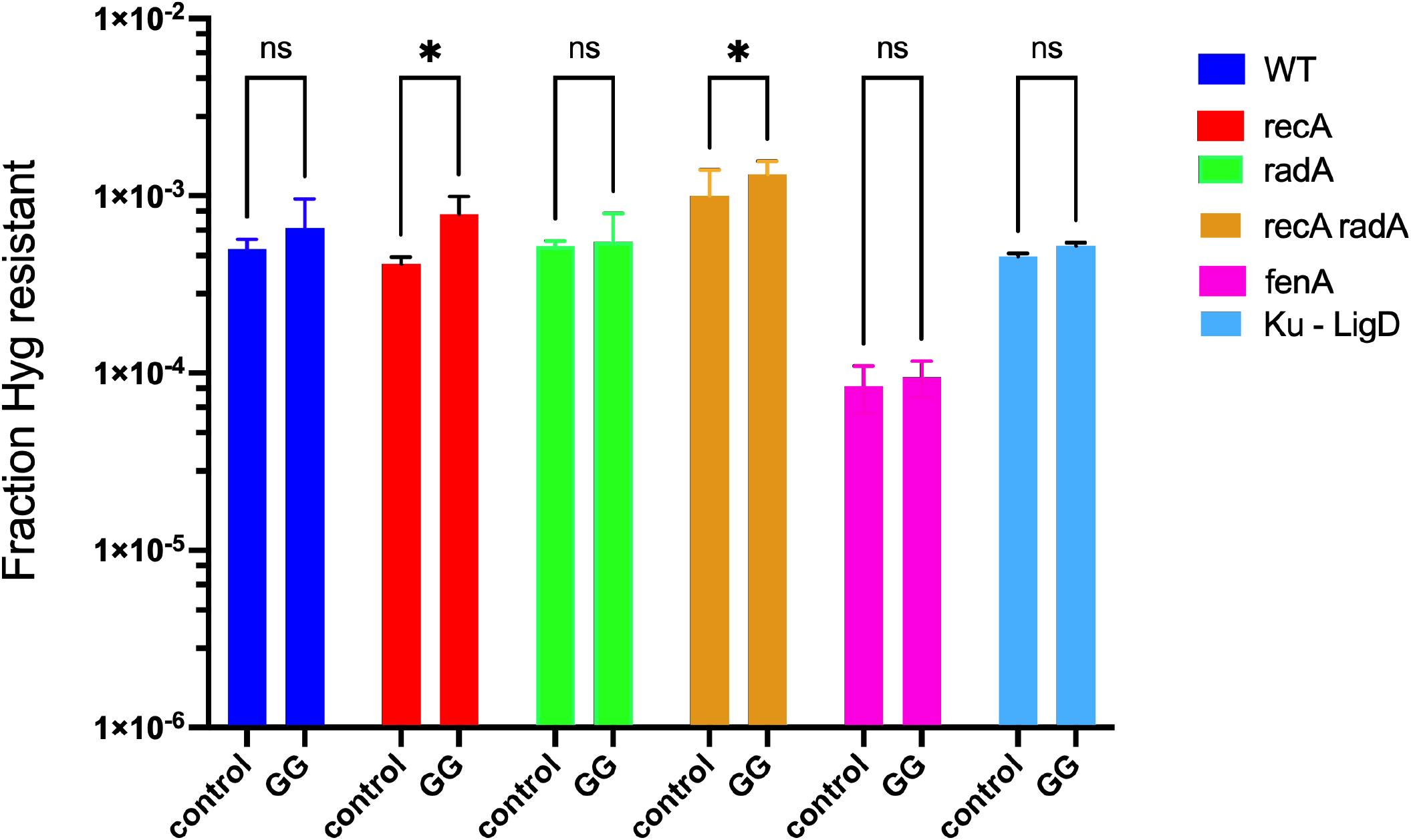
Testing of functions suspected in *M. smegmatis* NucS-promoted MMR repair. Oligos generating C-T mismatches that promote Hyg^R^ and generate no other mismatch (control), or a G-G mismatch 20 bp away from the C-T mismatch (G-G), were electroporated into *M. smegmatis* strain mc2-155 and a series of deletion mutants in genes suspected of being involved in *nucS*-promoted MMR. All strains contained the RecT-producing plasmid pKM461. Disruptions of genes involved in NucS-promoted MMR are expected to generate low frequencies of Hyg^R^ colonies with oligos promoting a repairable mismatch (G-G), compared to high frequencies of Hyg^R^ expected with oligos not generating the G-G mismatch (control oligo). Experiments were done in triplicate; standard deviations are shown. A 2-way ANOVA statistical analysis was performed comparing the frequencies of WT and each mutant, with and without the G-G mismatch; *t*-test, \**P* < 0.05.

This assay can also be used to identify genes that are involved in processing of a NucS-induced nick. The absence of a resident 5’-3’ exonuclease or helicase involved in processing would likely lead to interruption of the *hyg* reading frame by loss of base pairs other than 3 (or multiples of 3), again leading to loss of Hyg^R^ transformants relative to WT. Other functions in *M. smegmatis* were thus tested in this assay and included a strain mutant in both Ku and LigD (21,33), to see if NHEJ had a role in mycobacterial MMR, and the MSMEG_3883 gene which encodes *fenA*, a flap endonuclease involved in processing of 5’ overhangs (34,35). Results of this experiment are shown in Fig. 8 show that neither NHEJ functions LigD and Ku (MSMEG_5570 and MSMEG-5580) or the FenA flap endonuclease (MSMEG_3883) are involved in mycobacterial mismatch repair, as none of these functions lead to loss of Hyg^R^ transformants with the oligo that generates the G-G mismatch relative to the control oligo. The absolute frequencies of Hyg^R^ transformants is lower when performed in the *fenA* mutant strain relative to the other hosts, which is attributed to a possible effect on the frequency of oligo-mediated recombineering (which was not further addressed in this study).

Functions thought to be important players in NucS-promoted MMR in mycobacteria, such as other dsDNA exonucleases and helicases, are currently being examined using this double-mismatch oligo-recombineering assay.

## Discussion

The discovery of the EndoMS/NucS function, principally found in archaea and actinobacterial species, revealed that a new pathway for DNA mismatch repair had been identified that did not follow the nearly ubiquitous paradigm of MutS DNA mismatch recognition, followed by MutL (or MutH) nicking of the duplex strand containing the “wrong” base. Instead, in vitro characterizations of the NucS proteins from the archaeal species *T. kodakarensis* and the actinobacterium *C. glutamicum* revealed that while they recognized substrates containing mismatched substrates, these proteins proceed to make cuts in both strands in DNA mismatched-containing substrates. The cutting of dsDNA containing mismatches with *T. kodakarensis* NucS showed that the cut left five nucleotide 5’ overhangs and recessed 3’-OH ends leaving ligateable sticky ends characteristic of restriction enzymes. Further support of the dsDNA cutting capability of NucS comes from the Xray structural analysis of the NucS protein from the *T. kodakarensis* which clearly has structural characteristics of a restriction enzyme: a homodimeric structure consisting of two domains, an N-terminal domain acting as a dimerization function that recognizes the mismatched bases that are flipped out (like restriction enzyme Ecl18kl), and a C-terminal domain that contains the active site residues for cutting dsDNA in a manner reminiscent of type II restriction enzymes (14). These studies strongly suggest that NucS acting in archaea and actinobacterial species act on mismatched bases by creating a dsDNA break at the site of the mismatch, though this cutting has not been demonstrated in vivo.

A first step to examine the capability of NucS in mycobacteria to promote such a dsDNA break was investigated by Castenda-Garcia *et al* (19), who showed that an *M. smegmatis* Δ*nucS* strain showed a mutagenic phenotype when plated on rifampicin plates, which we confirmed here. However, their biochemical analysis revealed that while NucS bound to ssDNA, it did not bind to or cut substrates containing DNA mismatches, leaving open the possibility that NucS from mycobacteria might not behave in a similar manner to NucS from *C. glutamicum* (despite a 73% sequence identify between the two proteins). The most likely scenario for their failure to see cutting by *M. smegmatis* NucS was that, like NucS from *C. glutamicum,* it requires an interaction with the β-clamp. To this end, our ability to complement the mutagenic phenotype of *M. smegmatis* Δ*nucS* with full-length NucS, but not one missing the last 5 amino acids containing the PIP domain, is consistent with this supposition. Verification of this interaction between *M. smegmatis* NucS and its β-clamp will require biochemical analysis of these functions.

We then leveraged the oligo-mediated recombineering capabilities of *M. smegmatis* to deliver defined mismatches directly to the mycobacterial chromosome to examine the mismatch specificity of NucS MMR. We found that mismatches containing G-T, G-G, and T-T were specifically repaired using the NucS MMR pathway, while none of the other mismatches were corrected. The mismatch recognition specificities revealed here *in vivo* are consistent with previous specificities shown in *C. glutamicum* NucS_Cg_ in vitro. (17,18). They are also consistent with the mutational specificity of *M. smegmatis* reported by Castaneda-Garcia *et al* (24), who showed a high rate of both G:C > A:T and A:T > G:C transitions in *M. smegmatis nucS* strains (relative to wild type strains) using mutational accumulation experiments. This result agrees with the recognition of G-T mismatches by NucS in vivo, which would prevent the accumulation of transition mutations in wild type cells. However, while large increases in transversions are not observed in mycobacterial *nucS* mutant MA studies (3-fold increase at best relative to wild type (24)), mismatches of G-G and T-T that would lead to such transversion mutants are recognized and cut by mycobacteria NucS in vivo at essentially the same efficiency at G-T mismatches (this report). This discrepancy may be resolved by the supposition that G-T base pairs are by far the most highly generated mismatch generated in vivo, the repair of which has evolved to be quite efficient, and that G-G and T-T mismatches, while recognized and repaired by NucS, are simply not generated at the same high frequencies as G-T mismatches in vivo.

To unravel the mechanistic details of NucS MMR, we used oligo-mediated recombineering to examine exonucleolytic processing that takes place in the chromosome following the cutting of the chromosome at a repairable mismatch. This was done by delivering an oligo containing a G-G mismatch by recombineering and measuring either loss or retainment of a nearby mismatch (C-T) as determined by the frequency of Hyg^R^ colonies following plating of the outgrowth. The assay, however, cannot distinguish between whether a nick or a dsDNA cut is made at the site of the mismatch, as this recombineering approach can only measure the activity of endogenous exonucleases on the strand containing the mispaired bases supplied by the oligo (the top strand of the oligo highlighted in blue in Fig. 6B).

However, by comparing the number of bases needed to process that top strand by either a 5’ end or a 3’ end, it is clear that the 5’ end is processed more efficiently than the 3’ end. This can be seen by recognizing that the 5’ end in Fig. 7C needs 8 bases removed from the top strand before it reaches the T-C mismatch, which lowers the frequency of Hyg^R^ colonies (see results of the −5 oligo in Fig 7B). Significantly, this frequency is down nearly 100-fold relative to a 3’ end digestion of the top strand with mismatches generated by oligo +8 in Fig. 7B, which requires 6 base pairs to reach the C-T mismatch. Thus, 5’-3’ exonucleolytic processing is heavily favored at the site of a NucS-promoted nick or dsDNA break.

However, what is more telling in this experiment is the limited nature of the processing of the 5’-3’ exonuclease following the cut made by NucS at a mismatch. By examining the extent of loss of Hyg^R^ transformants among the oligos that generate mismatches that confer Hyg^R^ (C-T) and a substrate for NucS cutting (G-G), oligos “-5” to “-20” in Fig. 7B, one can observe that processing of the NucS-induced cut by a 5’-3’ exonuclease in most cases extends to 8 bases from the cut site (including the five base 5’ overhang). Digestion of an additional 3 bases (11 total) is diminished 20-fold, as evidenced by a 20-fold increase in the frequency of Hyg^R^ transformants seen with oligo “-8” relative to “-5” in Fig. 7B. Ultimately, the use of an oligo where the distance between the C-T and G-G mismatches generated at the replication fork is 20 bases (oligo “-20”) results in no loss of Hyg^R^ transformants relative to the control oligo. These results suggest that digestion of the 5’-3’ strand from the site of a NucS cut (or nick) involves at least 8 base pairs, but not more than 10-13 base pairs. This result means that if NucS makes a dsDNA cut at a mismatch in vivo, it is not further processed by RecA-promoted homologous recombination events, as efficient recombination requires much larger sequences of homology (∼200 bases) relative to what is observed in here (36).

The use of a double-mismatch generating oligo was also used to test the genetic dependency of NucS MMR. It is reasoned that if a NucS-promoted dsDNA cut is made in vivo at a mismatch, then inactivating mutations of any downstream functions would lead to non-repair of the dsDNA cut and subsequent death of the cell. In a similar vein, if only a nick is made at the mismatch by NucS, loss of any functions required for exonucleolytic processing of the nick would likely lead to some frequency of indel formation and thus to loss of Hyg^R^ transformants. Our results showed that neither RecA nor RadA is involved in downstream processing of the cut made by NucS, consistent with the limited processing of cuts made in vivo shown in Fig. 7. We next examined if the non-homologous end-joining functions Ku and LigD might somehow be involved MMR in *M. smegmatis*. Since mycobacteria exhibit the uniqueness of possessing both the novel NucS MMR system and NHEJ functions (neither present in most bacteria), it was hypothesized that NHEJ, together with restriction enzyme-like NucS, might work together to repair mismatches following a NucS-promoted dsDNA cut, in a way the exhibits a high-fidelity version of NHEJ-promoted repair of dsDNA breaks. However, we found no effect on the frequency of Hyg^R^ transformants with the double-mismatch generating oligo in a strain deleted of both Ku and LigD, showing that NHEJ functions are not involved in MMR in mycobacteria. Finally, the limited processing of the cut made in vivo was highly suggestive of a flap endonuclease, where a cut is made at the ssDNA-dsDNA junction to remove a 5’ single-stranded DNA flap. In this scenario, one imagines that NucS cutting in vivo is limited to only the replicative strand (by some unknown function) and that unwinding from the nick is limited to the generation of a 5’ flap containing the mispaired base. Subsequent cutting of the flap by a resident endonuclease was suspected. However, the deletion of the gene that encodes the FenA 5’-flap endonuclease (MSMEG_3883) had no effect on the frequency of Hyg^R^ transformants in our double-mismatch generating oligo assay, ruling out a role of *fenA* in NucS-promoted MMR, or that some other exonuclease in vivo can substitute for FenA in its absence. In these types of cases, it may require double (or multiple) exonucleolytic mutants to be generated to determine if processing of the NucS-induced break is carried out by FenA-like overlapping functions.

Thus, downstream components of the NucS MMR pathway remain elusive as we continue to use this assay to examine the role of other nucleases and helicases known to exist in the mycobacterial genome that may have a role in high fidelity repair of mismatched bases. If in fact, the mycobacterial NucS protein makes a dsDNA break at the mismatch in vitro (like its *C. glutamicum* counterpart), the mechanism in vivo would have to rely on a modification of NucS activity in vivo (perhaps by binding of an interacting partner) such that nicking of the leading strand is prevented. Alternatively, one can imagine that a dsDNA cut at the mismatch is made in vivo but is quickly held together via the action of an SMC-like protein (perhaps the *rad50* analog MSMEG 5590) that promotes a scaffold for quick religation of the leading strand that doesn’t expose the cell to the deleterious effects of a dsDNA break. Further strains containing deletions of such genes will be constructed and tested in our double-mismatched oligo assay.

## ACKNOWLEDGEMENTS

We wish to acknowledge Michael Glickman for the *recA* knockout strain and the *ku-ligD* double mutant. We thank Christopher Sassetti and Kadamba Papavinasasundaram for helpful discussions, materials support and managerial assistance, and Peter Oluoch and Rick Baker for help with statistical analyses.

## FUNDING

This work was supported in part by the Bill and Melinda Gates Foundation grant # INV-040487.

## CONFLICT OF INTEREST

Authors declare no conflicts of interest.

## Notes

### Competing Interest Statement

The authors have declared no competing interest.

### Summary of Updates

Typos corrected; sections of text revised for clarity; replicate data added to Figures 7 and 8; Figure S1 axes clarified.

