## Supplementary Tables and Figures for "*Mycobacterium smegmatis* NucS-promoted DNA mismatch repair involves limited resection by a 5’-3’ exonuclease and is independent of homologous recombination and NHEJ"

A.

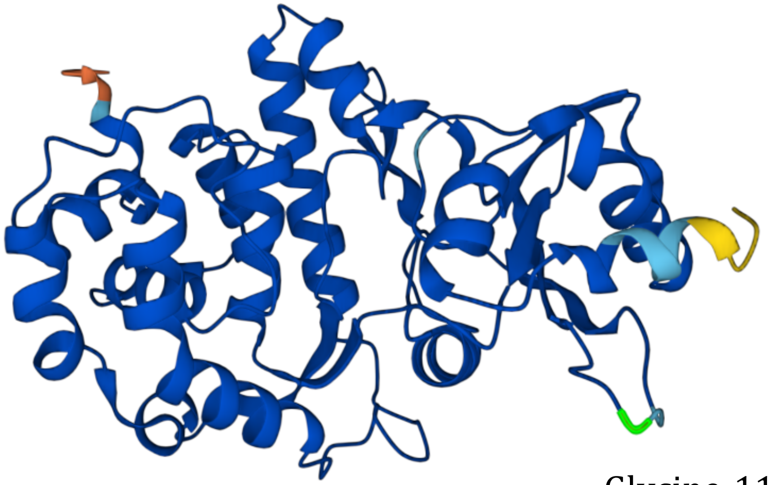

Glycine-110

B.

| | |
|---|---|
| *Streptomyces hygroscopicus1* | GRGELRPG**TGA**WPWPYLV |
| *Streptomyces varsoviensis* | GRGELRPG**GAG**WPWPYLV |
| *Streptomyces griseocarneus* | **A**RGELRPGT**D**AWPWPFLV |
| *Streptomyces luteoverticillatus* | GRGEL**F**PG**GDG**WPWPYLV |
| *Saccharothrix australiensis* | G**A**GELRPG**SPT**WPWPFLV |
| *Streptoalloteichus hindustanu* | GRGELRP**SGPG**WPWP**YV**V |
| *Streptomyces hygroscopicus* | GRGELRPGT**R**AWPWPYLV |

Figure S1

### Figure S1

**Description of the defective hygromycin phosphotransferase gene.** (A) In order to develop an assay where one could generate multiple types of mismatches at a stop codon placed within a selectable gene, which would also require the protein to still be active independent of the type of amino acid replacing the stop codon, we turned to the gene that confers hygromycin resistance ($Hyg^R$). In the alpha fold structure of the hygromycin phosphotransferase gene, an extended loop at the surface of the protein was found to include a glycine at position 110. We thus replaced the glycine codon GAA with the TGA stop codon using recombineering. The assay ultimately revealed that all amino acid substitutions made at this position were tolerant.

(B) The capability of replacing glycine-110 in *hyg* with multiple types of amino acids was also favored following a search of the protein sequences from various hygromycin phosphotransferases in GenBank. Clearly, Gly-100 is not highly conserved.

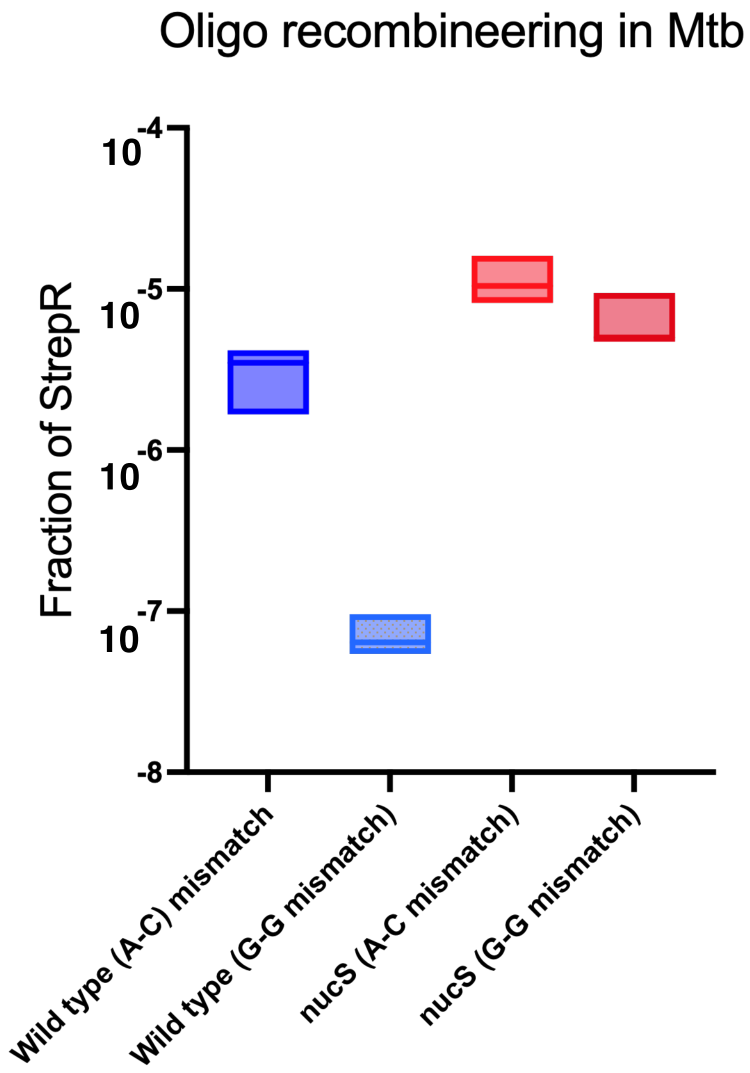
Oligo recombineering in Mtb
Fraction of StrepR
$10^{-4}$
$10^{-5}$
$10^{-6}$
$10^{-7}$
-8
Wild type (A-C) mismatch
Wild type (G-G mismatch)
nucS (A-C mismatch)
nucS (G-G mismatch)

Figure S2

### Figure S2

**Recombineering of the *rpsL* locus in *M. tuberculosis*.** An assay was set up like the one shown in Table 1 but testing MMR in *M. tuberculosis*. Both wild type (H37Rv) and a ΔnucS derivative (ΔRv3122) were electroporated with oligos targeting the *rpsL* gene, where either a predicted repairable mismatch (G-G) or unrepairable mismatch (A-C) was generated. In H37Rv cells, the A-C mismatch is left unrepaired leading to high levels of streptomycin resistance, whereas the G-G mismatch is repaired leading to low levels of streptomycin resistance. Both mismatches lead to high levels of streptomycin resistance in the ΔnucS derivative. This experiment was caried out in a BSL3 facility governed by the Institutional Biosafety Office at UMass Chan Medical School.

### Table S1

### Hyg repair oligos used in the mismatch specificity assay

| | | Hyg repair oligos | mismatch | new codon | Amino acid |
|---|---|---|---|---|---|
| Wild type | 5’ | ATCCGGCTCATCACCAGGTAGGGCCACGGCCAGGC**TCC**GGTGCCGGGCCGCAGCTCGCCGCGGCCGAGGA 3’ | | | |
| Stop codon | | ATCCGGCTCATCACCAGGTAGGGCCACGGCCAGGC**CTA**GTTGCCGGGCCGCAGCTCGCCGCGGCCGAGGA | | | |
| Oligo # | | Position (1) 110 GLYCINE G | | | |
| 1 | | ATCCGGCTCATCACCAGGTAGGGCCACGGCCAGGC**TCC**GGTGCCGGGCCGCAGCTCGCCGCGGCCGAGGA | multiple | GGA | glycine |
| 2 | | ATCCGGCTCATCACCAGGTAGGGCCACGGCCAGGC**CTT**GGTGCCGGGCCGCAGCTCGCCGCGGCCGAGGA | T/T | AAG | Lysine |
| 3 | | ATCCGGCTCATCACCAGGTAGGGCCACGGCCAGGC**CTG**GGTGCCGGGCCGCAGCTCGCCGCGGCCGAGGA | G/T | CAG | Glutamine |
| 4 | | ATCCGGCTCATCACCAGGTAGGGCCACGGCCAGGC**CTC**GGTGCCGGGCCGCAGCTCGCCGCGGCCGAGGA | C/T | GAG | Glutamic Acid |
| | | Position (2) 110 GLYCINE G | | | |
| 5 | | ATCCGGCTCATCACCAGGTAGGGCCACGGCCAGGC**CGA**GGTGCCGGGCCGCAGCTCGCCGCGGCCGAGGA | G/A | TCG | Serine |
| 6 | | ATCCGGCTCATCACCAGGTAGGGCCACGGCCAGGC**CCA**GGTGCCGGGCCGCAGCTCGCCGCGGCCGAGGA | C/A | TGG | Tryptophan |
| 7 | | ATCCGGCTCATCACCAGGTAGGGCCACGGCCAGGC**CAA**GGTGCCGGGCCGCAGCTCGCCGCGGCCGAGGA | A/A | TTG | Leucine |
| | | Position (3) 110 GLYCINE G | | | |
| 8 | | ATCCGGCTCATCACCAGGTAGGGCCACGGCCAGGC**GTA**GGTGCCGGGCCGCAGCTCGCCGCGGCCGAGGA | G/G | TAC | Tyrosine |
| 9 | | ATCCGGCTCATCACCAGGTAGGGCCACGGCCAGGC**ATA**GGTGCCGGGCCGCAGCTCGCCGCGGCCGAGGA | A/G | TAT | Tyrosine |

The top two rows show the reverse complement of a central region of the *hyg* gene, with the position of the Gly codon and the stop codon in bold face.

### Table S2

#### Oligonucleotides used for ORBIT-generated mutations and Oligo-mediated recombineering

<u>Identifier (Gene) ORBIT integration plasmid</u>

**ORBIT:**

ΔMSMEG_4923 (ΔnucS) pKM464
TCAGAAGAGCCGGTACTCGTCGCTGTCCATTCCGCGCATCTGGTCGTAATCGAGTGTCACACAACGGATTGGTTTGT
ACCGTACACCACTGAGACCGCGGTGGTTGACCAGACAAACCGGGCAGAGGGCAGATGAGCGGTGAGCCGGCCGACGT
AGTCGACGGTGCACTGGGCTATCACGAGGCGCAC

ΔRv1321 (ΔnucS) pKM488
TTTGCGGCGCCGAGCCATCGCATCAGTTTAATCGCGCAACTCAGAACAGCCGGTACTCGCCGCTATCCATGGTTTGT
ACCGTACACCACTGAGACCGCGGTGGTTGACCAGACAAACCCTGGGCGATGACTAGACGCACCCGACTCACCTTAGA
GCGCGCAACGACGTTGTTCCTTAGAGCGTGACCG

ΔMSMEG_6079 (ΔradA) (pKM611)
CGCAGGCCCGGCTGTCGGTGCCTGCCGATACTGTCACGGCGTGGCCGGTTCGAAAATACGTTCGCAGTACGGTTTGT
CTGGTCAACCACCGCGGTCTCAGTGGTGTACGGTACAAACCCGGGAGATCGCGATTGCGGGTGCCCAATAGGATCGG
GACAATACCGGCCGATCCTAGGAGGGCTGATGGC

ΔMSMEG_3883 (ΔfenA) deletion (pKM611)
GTCATGATTCACGATACGTGGGACCGCGACTGTCCCTGCCGTGGTGTGTGACATCGACGTCGAGGTAGCCGGTTTGT
CTGGTCAACCACCGCGGTCTCAGTGGTGTACGGTACAAACCCAGACCGCACTGGACCAGCTGCCCGACTGAGCCGGTC
TACTTCGGCCGGCCGACCTCGTAGGTGCCGTCG

MSMEG_1397- rpsL intergenic region insertion (pKM614)
CACCACAATACCAGGGCCGACCGCGACGAAACAAACTCGTGCACGGCGGTCTAATCGCAGGTCAGACAGTGGTTTGT
ACCGTACACCACTGAGACCGCGGTGGTTGACCAGACAAACCGAGGTCCACCCTACGCTTCGCGACGCTCCATCCCGGA
GGCCAGGCGCAGCAGCATATCGGTGAAAACCGC

**Oligo-Recombineering:**

rpsL-K43R (base substitution)
CGCGGGCAACCTTCCGAAGCGCCGAGTTCGGCTTCCTCGGAGTGGTGGTGTACACGCGGGTGCATACACC

rpsL-K43N (base substitution)
CGCGGGCAACCTTCCGAAGCGCCGAGTTCGGCTTGTTCGGAGTGGTGGTGTACACGCGGGTGCATACACC

Smeg-leuB-R101+1 (one bp insertion)
AGCGGGCCCGTCACACCCGGATACAGCCTGCCGGGGCGGCAGATTCACGTGATGGTCGAGCGCGAACCGCAGTT

Smeg-leuB-R101Δ1 (one bp deletion)
AGCGGGCCCGTCACACCCGGATACAGCCTGCCGGGGCCAGATTCACGTGATGGTCGAGCGCGAACCGCAGTT

Smeg-leuB-R101Δ2 (two bp deletion)
AGCGGGCCCGTCACACCCGGATACAGCCTGCCGGGGCAGATTCACGTGATGGTCGAGCGCGAACCGCAGTT

Smeg-leuB-WT
AGCGGGCCCGTCACACCCGGATACAGCCTGCCGGGGCGCAGATTCACGTGATGGTCGAGCGCGAACCGCAGTT

Smeg-NucS-D138A
GCGACCGAACGGCCCAGCTCGTCGCGGCACAACAGGGCGACGGGCCCGATCGGGGTCGGATACTCGCGGCGCA

Smeg-NucS-E152A
TGTTCGACGCCGTCGATCTCGCCGCGGCGCTTGATCGCCACGGCGACCGAACGGCCCAGCTCGTCGCGGCACA

Smeg-NucS-K154A
GTCAGCTGTTCGACGCCGTCGATCTCGCCGCGGCGCGATCTCCACGGCGACCGAACGGCCCAGCTCGTCGC

Hyg repair
GACGCCCCGGTGCCGGTGCCCCGCCTCCTCGGCCGCGGCGAGCTGCGGCCCGGCACCTcGGCCTGGCCGTGGCCCTACC
TGGTG

Hyg repair-32
GACGCCCCGGTGCCGGTGCCCCGCCTgCTCGGCCGCGGCGAGCTGCGGCCCGGCACCTcGGCCTGGCCGTGGCCCTACC
TGGTG
